# Forecasting crowd dynamics through coarse-grained data analysis

**DOI:** 10.1101/175760

**Authors:** Sebastien Motsch, Mehdi Moussaïd, Elsa G. Guillot, Mathieu Moreau, Julien Pettré, Guy Theraulaz, Cécile Appert-Rolland, Pierre Degond

## Abstract

Understanding and predicting the collective behaviour of crowds is essential to improve the efficiency of pedestrian flows in urban areas and minimize the risks of accidents at mass events. We advocate for the development of a & “crowd forecasting system„whereby real-time observations of crowds are coupled to fast and reliable models to produce rapid predictions of the crowd movement and eventually help crowd managers choose between tailored optimization strategies. Here, we propose a Bi-directional Macroscopic (BM) model as the core of such a system. Its key input is the fundamental diagram for bi-directional flows, i.e. the relation between the pedestrian fluxes and densities. We design and run a laboratory experiments involving a total of 119 participants walking in opposite directions in a circular corridor and show that the model is able to accurately capture the experimental data in a typical crowd forecasting situation. Finally, we propose a simple segregation strategy for enhancing the traffic efficiency, and use the BM model to determine the conditions under which this strategy would be beneficial. The BM model, therefore, could serve as a building block to develop on the fly prediction of crowd movements and help deploying real-time crowd optimization strategies.

## 1 Introduction

Since 2008, more than half of the world population is living in cities [1], which asks for the development of large-scale multimodal transportation systems. At the core of such systems, crowds of pedestrians are to be found in transportation facilities. The massive presence of this population in airports, train stations, metro stations, etc. is a challenge in terms of safety and transportation efficiency [2]. This paper deals with the experimental analysis and modeling of crowd movements so as to develop operational tools to manage pedestrians in places as various as airports, train or metro stations or streets.

Managing the comfort and safety for pedestrians is difficult first because they are not operated or regulated as simply as vehicles can be through sets of rules and traffic signals [3]. Hence, it is difficult to prevent the formation of dense traffic areas, where compacted crowds are exposed to risks of stampedes [4, 5]. For this reason, it is important to design pedestrian traffic management systems similar to those existing for car traffic, and more particularly simulation tools to predict the short-term evolution of the traffic from conditions measure *in-situ* and in real time.

Today, on the one hand, pedestrian crowd simulators are developed for the purpose of validating the layout and the structure of buildings aimed at hosting a large public.This validation is performed beforehand, at the stage of design [6]. Most are not adapted to an on-line usage.Nevertheless, on the other hand, recent progress in pedestrian tracking technologies, both for detailed trajectories [7, 8, 9] or global traffic conditions [10, 11, 12], make possible to initialize a pedestrian traffic simulators with current conditions and to perform short-term traffic predictions. This would open the possibility to anticipate risks of congestion and to react accordingly. Such simulators are lacking today, thus, it is important to focus our attention on simulators which are able to continuously process a flow of input traffic data and which are efficient enough to perform short term predictions on-line. Our work aims at to fill this particular gap, by developing online simulation tools to assist the management of pedestrian traffic.

We aim at performing short term predictions in places presenting risks of large attendance and risk of congestion. A specific type of environment retains our attention: corridors. By corridors, we mean elongated areas such as metro corridors, sidewalks or shopping arcade where pedestrians form bidirectional traffic flows. This type of place is particularly interesting to study because corridors generally link large places with high attendance (e.g., a corridor between two platforms). Because the traffic intensity is higher in corridors, they are the place where congestion can more easily initiate. In addition, because corridors are often organized in networks, they offer an opportunity to apply rerouting strategies to locally limit the risk of congestion: pedestrian would be suggested or forced to avoid corridors presenting a risk of future congestion, similarly to car drivers with modern GPS-based assistance systems.

How to perform accurate and fast predictive simulation for bidirectional pedestrian traffic in corridors? Two categories of models are described in the literature: microscopic and macroscopic approaches. The former approach is an ascending one: the motion of individual pedestrians is independently simulated based on multi-agent technique and crowd traffic conditions emerge from combination of interactions between agents. They are fundamentally based on the local models of interactions, which were developed in various disciplines such as physics [13], computer graphics [14] or behavioral and cognitive sciences [15, 16], and which dictate how agents influence each other‚s motion. The latter type of approach are macroscopic ones. They model a crowd as a whole, a matter moving like a compressible fluid. Relying on the conservation law, a macroscopic simulation computes the crowd motion by estimating the temporal variations of local density [19]. Algorithmic complexity is one fundamental difference between these two types of simulators. Microscopic simulators consider interactions between all possible pairs of agent and are quadratic by nature. In contrast, macroscopic algorithms are linearly complex with the size of the crowd. This difference makes macroscopic approaches a model of choice. While microscopic models fail to produce real time simulations as soon as the pedestrian number becomes large, macroscopic models can provide fast, simple and yet surprisingly accurate results even at large densities.

In this paper, we propose a macroscopic model to predict bi-directional traffic in corridors. To enhance computational efficiency, we consider only the longitudinal direction of the walkway, ignoring the lateral dimension. We show hereafter that despite this simplification a macroscopic model is able to quantitatively reproduce key features of crowd dynamics. In such category of model, the relevant information encoding crowd dynamics is the fundamental diagram, which that defines pedestrian flux as a function of pedestrian densities. More precisely, the proposed method relies on a “Bi-directional Fundamental Diagram” (BFD), which captures situations of a crowd made of people moving in the same and opposite directions. Fundamental diagrams are widely used for one-way traffic (since the pioneering work of Lighthill and Whitham [20]) but two-way pedestrian traffic has been scarcely investigated under the angle of the BFD [21, 22, 23, 24, 25, 26, 27]. Yet, two-way traffic is the most common situation in everyday life and counter-flows played a decisive role in several crowd disasters [28, 29].

To characterize the BFD, several experiments have been conducted from which we establish the first result of the present paper: an estimation of the BFD from experimental data. The data are acquired in real time tracking experiment using automatic motion capture techniques [30]. Our second result is the validation of the Bi-directional Macroscopic (BM) model which uses the calibrated BFD as its core. Comparisons between simulations and experiments demonstrate that the BM model captures essential features of crowd dynamics. We then test our model in a typical forecasting scenario where the crowd is recorded at two distant points. We show that the model accurately predicts the state of the corridor between the recording points and develops traveling waves similar to those observed in the experiments. Our third result is the determination of the optimum of a corridor segregation strategy consisting in separating or not antagonist pedestrian fluxes according to the corridor occupancy. The BM model suggests that the segregation is efficient when the two antagonist fluxes are roughly balanced, but is counter-productive in strongly unbalanced cases.

## 2 Methods

### 2.1 Experimental setup

In this section, we describe our experiment on pedestrian traffic. Our objective was to build a Bi-directional Fundamental Diagram (BFD), i.e., to observe the relation between the moving speed of participants under different conditions of density and balance of counter-flows. To this end, we asked participants to walk in a circular corridor delimited by walls (as displayed in Figure 1a). Some participants were instructed to walk in the clockwise direction, some others to walk in the anticlockwise direction. They were assigned a walking direction before each trial. For each trial, participants were initially still in the corridor and their positions randomly distributed. Each replication lasted 60 seconds after starting signal was given.

**Figure 1:**
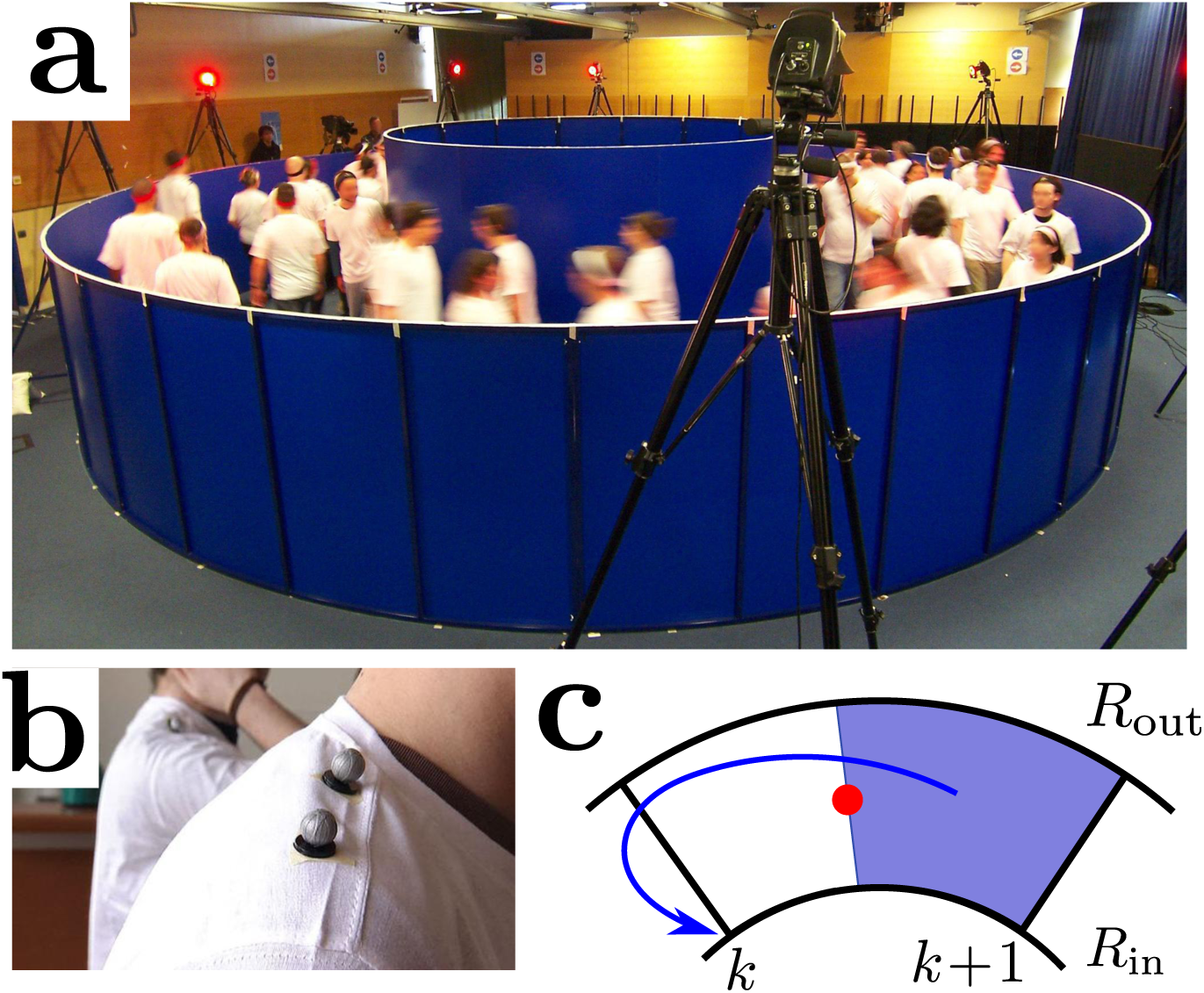
Experiments and data acquisition. (**a**) A typical experiment where bidirectional circulation is analyzed. (**b**) Participants equipped with reflexive markers were tracked by means of an optoelectronic motion capture system (VICON MX-40, Oxford Metrics, UK). (**c**) The area-weighting assignment procedure: The particle *i* (red dot) is located in the cell 𝒞^*n*^(*i*) = [*θ*_*k*_*, θ*_*k*+1_] and is assigned to the two nodes *θ*_*k*_ and *θ* in proportion to the area enclosing the opposite node. For instance, on the picture, the assignment to the node *θ* is in proportion to the ratio of the shaded area to the area of 𝒞^*n*^(*i*).

Participants were not allowed to communicate, and were asked to behave as if they were walking alone in a street to reach a destination. 119 volunteers participated the experiments (we performed two experimental sessions, one with 59 participants and one with 60 participants). They were adults recruited through advertising and with no known pathology which would affect their locomotion. The experiment conformed to the declaration of Helsinki. Participants were naive with respect to the purpose of our experiments.

Experiments took place in 2009 in Rennes, France. The circular corridor internal and external radius was respectively 2 and 4.5 meters. We studied 3 different proportions of fluxes, i.e., the balance of participants walking in a direction versus participants walking in the opposite direction. We studied: 100%–0% balance, 75%–25% balance and 50%–50% balance.

We collected kinematics data. To this end, participants wore white T-shirts and 4 reflexive markers (see Fig.1b), one on the forehead, one on the left shoulder, and two on the right shoulder. This made the distinction of the left and right shoulders easier.The motion of the markers has been tracked by 8 infra red cameras placed all around the experimental setting. Marker trajectories have been reconstructed using Vicon IQ software (VICON MX-40, Oxford Metrics, UK). The center of mass of the 4 markers projected onto the horizontal plane is computed and recorded as the position of the subjects [30, 17].

### 2.2 Data processing

The collected data give access to the two-dimensional Cartesian coordinates (*x*_*i*_*, y*_*i*_)(*t*^*n*^) (relative to the center of the arena and a fixed reference frame) of each pedestrian, as well as their associated polar coordinates (*θ*_*i*_*, r*_*i*_)(*t*^*n*^) (with *x*_*i*_ = *r*_*i*_ cos *θ*_*i*_, *y*_*i*_ = *r*_*i*_ sin *θ*_*i*_), sampled at a frequency of 10 Hz (i.e. *t*^*n*^ = *n* Δ*t* with Δ*t* = 0.1 s). Since we are interested in a one dimensional analysis, we focus on the angular coordinate of the pedestrians. We assign a fixed value for the radial coordinates *R*_med_ = 3.25*m* corresponding to the median value. To estimate the density distribution of pedestrians on the arena, we use a classical area-weighting procedure (a.k.a. Particle-In-Cell method [54, 55]). This procedure consists in assigning each pedestrian to the two neighboring nodes in proportion to the length enclosing the opposite node (see Fig. 1c).The density 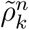 is obtained by combining the contributions of all pedestrians *i* belonging to the two cells sharing the node *k*, according to the formula

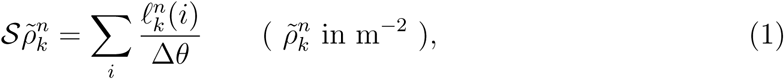

where the sum extends over pedestrians *i* belonging to one of the two cells sharing node *k*. The quantity *S* is the area of the cross section

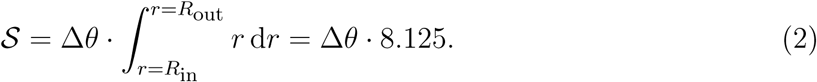

By linear interpolation, this procedure gives rise to a continuous piecewise linear reconstructed density 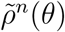 at time *t*^*n*^. We can verify that the total number of pedestrian *N* is preserved in this interpolation procedure, i.e 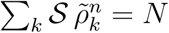.

To estimate the flux, we compute a finite-difference approximation of the azimuthalcomponent of the velocity 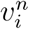 of each pedestrian i at time *t*_*n*_:

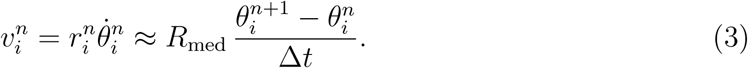

The densities 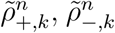 and fluxes 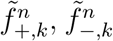 of the clockwise and anti-clockwise pedestrians respectively are estimated at time *t*^*n*^ on the polar grid *θ*_*k*_ = *k*Δ*θ* by means of the area-weighting assignment procedure described in Fig. 1c.The procedure for estimating the densities 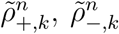 uses Eq. 1 where the sum is restricted to pedestrians moving in the considered direction.For the estimation of the fluxes, we use the following formula

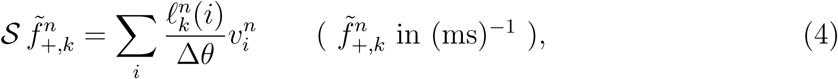

where again the sum extends over pedestrians *i* belonging to one of the two cells sharing node *k* (and moving in the considered direction).

### 2.3 Model

To investigate the flow of the pedestrians and make prediction, we consider a bidirectional extension of the traffic model [20, 24]. Let *ρ*_*±*_(*x, t*) be the densities at position *x* (along the median circle) and time *t*. The Bi-directional Macroscopic (BM) is written:

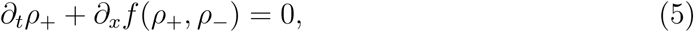

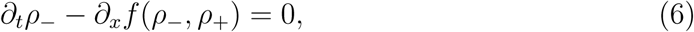

where *f* (*ρ*_+_*, ρ*_−_) is the “Bi-directional Fundamental Diagram”(BFD) (see Fig. 2a). Both Eq. 5 and Eq. 6 have the form of a continuity equation for the associated pedestrian density. It expresses that the only cause of a time variation of any of these densities in a corridor stretch [*x, x* + d*x*] is due to the flux of pedestrians entering this stretch at *x* and leaving it at *x* + d*x* (or vice versa according to the direction of motion). Because *x* corresponds to the angular coordinate *θ*, *ρ*_*±*_ are assumed periodic with period 2*πR*_med_. Since the model is a system of conservation laws, we use a finite volume method to estimate numerically its solutions [56, 57]. More specifically, we use a central scheme method [24, 58]. By analogy with the one-directional traffic model, we introduce characteristic speeds of the models (see Fig. 2b):

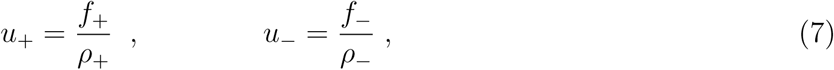

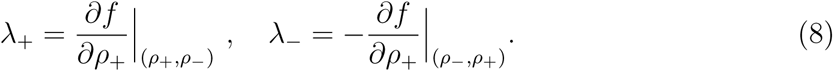

**Figure 2:**
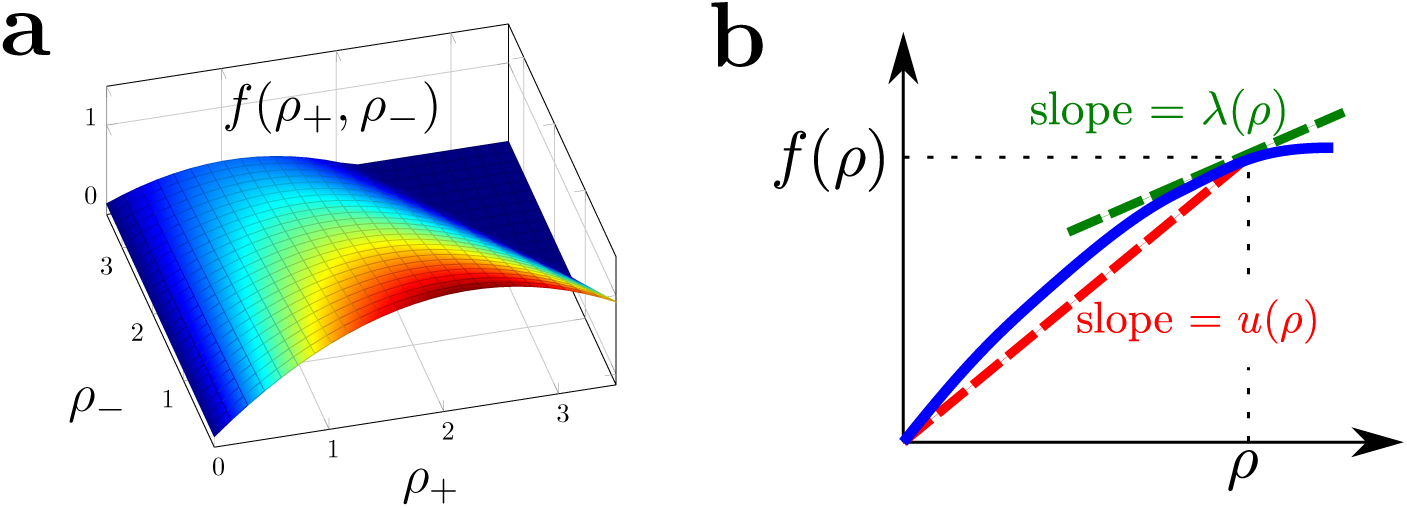
Characteristic speeds in the BM model. (**a**) Perspective view of the BFD *f* (*ρ*_+_*, ρ*_−_) as a function of *ρ*_+_ and *ρ*_−_. (**b**) Sketchy fundamental diagram (in red) of a one-way flow *f* (*ρ*) as a function of the single density *ρ*. The function *f* (*ρ*) has the same monotony as cuts of the BFD *f* (*ρ*_+_*, ρ*_−_) along lines *ρ*_−_ = Constant. The average velocity *u*(*ρ*) = *f*(*ρ*)/*ρ* and the cluster velocity *λ*(*ρ*) = *f*′(*ρ*) are respectively the slopes of thesecant (in red) and tangent (in green) lines to this curve.

Strictly speaking, the quantities *λ*_*±*_ are the velocities at which level curves of the densities *ρ*_*±*_ are convected. It corresponds to the velocity of waves that arise from a perturbation of a uniform solution given by *ρ*_*±*_. Since the model is actually bidirectional, the expression of *λ*_*±*_ in Eq.8 is actually an approximation (see appendix A).

## 3 Results

### 3.1. Estimated bi-directional fundamental diagram

From these data, we estimate the relation between the local densities *ρ*_+_ and *ρ*_−_ and fluxes of the clockwise and anti-clockwise pedestrians (respectively denoted by *f*_+_ and *f*_−_). Using the reconstructed densities and fluxes, to each couple (*k, n*), we can associate two triples 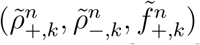 and 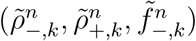 Thus, this provides us a sampling of *f*_+_ and *f*_−_. Toestimate the relation between densities and fluxes, we partition the (*ρ*_+_*, ρ*_−_) space intosquared sampling cells of size Δ*ρ* = 0.1 m^−2^. For each of these cells, we compute the average of the samples 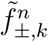 conditioned by the fact that 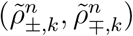 lies in the chosen cell. As a result, we obtain an estimated *f* (*ρ*_+_*, ρ*_−_) relationship. We then seek for a parametric estimation of the fundamental diagram *f* (*ρ*_+_*, ρ*_−_). The procedure consists in finding them best fit of the samples 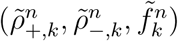 to a prescribed analytic formula. After several trials, it appears that the best fit consists in taking the velocity *u*_+_ = *f*_+_*/ρ*_+_ equal to an affine function of (*ρ*_+_*, ρ*_−_) leading to the formula (9). Different parametric estimations of the fundamental diagram are then performed according to the experimental conditions. This is needed because the subjects themselves or their physical or psychological conditions differ from one experimental trial to another one. We gather replications corresponding to the same flux balance and realize three different estimations for the 50%–50% balanced fluxes case, the 75%–25% flux balance case and the mono-directional (100%–0% flux balance) case. The result of the parametric estimation is given in Table 1 and the flux for the balanced case is represented in Fig. 2a.

In Fig. 3a, we plot the BFD for the experimental data corresponding to balanced fluxes (50% 50% balance). The grey area corresponds to the absence of data, since the observed maximum pedestrian density achieved in our experiments is about 2 m^−2^. It appears that the BFD is increasing in *ρ*_+_ and decreasing in *ρ*_−_ and reaches its maximum at the total density of 1.7 m^−2^.The estimation of the decay of the BFD as a function of *ρ*_−_ is one of the key results of this work. As *ρ*_−_ increases, the density of obstacles impeding the motion of the pedestrians intensifies, resulting in an increased ‚friction‚ [31]. We parametrize the flux f using a quadratic function of the form:

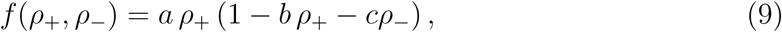

with densities expressed in (pedestrians) m^−2^ and flux expressed in (ms)^−1^. This parametric BFD is the key input in the Bi-directional Macroscopic model (see Methods).Despiteits simplicity, the formula is able to capture several key features of the pedestrian dynamics as we will see below. Coefficients *a*, *b* and *c* are easily interpreted and thus their estimation gives insights into the pedestrian dynamics. For instance, coefficient *a* corresponds to the free average velocity of the pedestrians. Coefficients *b* and *c* correspond to ‚friction‚ with respectively coand counter-moving pedestrians. Using linear regression, we obtain the estimated values given in table 1. As an example, the estimated flux for the experiments with 50% – 50% balance is plotted in Fig. 3b.

**Table 1:** Parametric estimation of the bi-directional fundamental diagram. The coefficients *a*, *b* and *c* refer to Eq. 9. To measure the accuracy of each regression, we estimate the coefficient of determination *R*^2^ in the last column (the closer *R*^2^ is to 1, the better the estimation is).

| Sample set | a | b | c | $R^2$ |
| --- | --- | --- | --- | --- |
| 50% – 50% | 1.218 | 0.273 | 0.181 | 0.944 |
| 75% – 25% | 1.216 | 0.087 | 0.203 | 0.972 |
| 100% – 0% | 1.269 | 0.077 | 0 | 0.982 |

**Figure 3:**
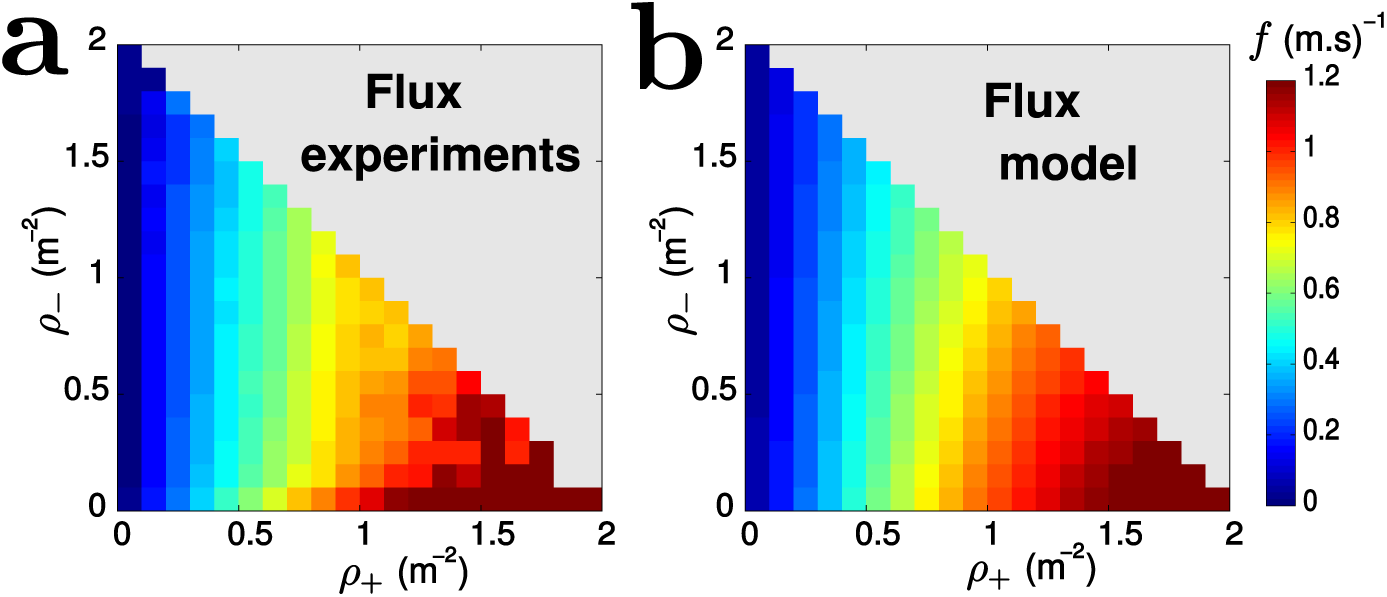
Bi-directional Fundamental Diagram (BFD). (**a**) Estimated BFD expressing the flux *f* in one direction as a function of the density of pedestrians moving in the same direction *ρ*_+_ and in the opposite direction *ρ*_−_. (**b**) Parametrized BFD used in the model. The values of *f*, in (*m.s*)^−1^, are color coded according to the color bar.

### 3.2 Data-model comparison: cluster dynamics

In order to test the model, we envision a crowd forecasting system where sensors are placed at two distant locations *A* and *B > A* of a corridor. The sensors record the densities and mean velocities of the incoming pedestrians. The model is then used to predict the occupancy of the corridor [*A, B*] knowing these boundary conditions. To match this situation with the ring-shaped corridor used in the experiments, we introduce a branch cut along the half line *θ* = 0 in polar coordinates and assume that *A* corresponds to *θ* = 0 and *B* to *θ* = 2*π*. Pedestrians crossing the branch cut in the clockwise (resp. anti-clockwise) direction are therefore entering the corridor at *B* and moving towards the left (resp. at *A* and moving to the right) as depicted in Fig. 4a. We use the densities estimated from the experiments as boundary and initial conditions for the BM model (see Methods section for a detailed description of the model and simulations). Fig. 4b,c (left) show the densities *ρ*_*±*_(*x, t*) for one of the experimental replications corresponding to 50%–50% flux balance with 60 pedestrians, as functions of *x* (horizontal axis) and *t* (vertical axis, running downwards) in color code, from blue (low density) to red (high density). Fig. 4b,c (right) show the results of the BM model run in the crowd forecasting situation as described above. Both data and simulations exhibit the formation of clusters (which appear in intense red colors on Fig. 4b,c), separated by almost vacuum regions (in intense blue color on the figure). Clusters move along with the pedestrians but not at the same speed. Similar comparisons for the 75%–25% flux balance are presented in figure 5.

**Figure 4:**
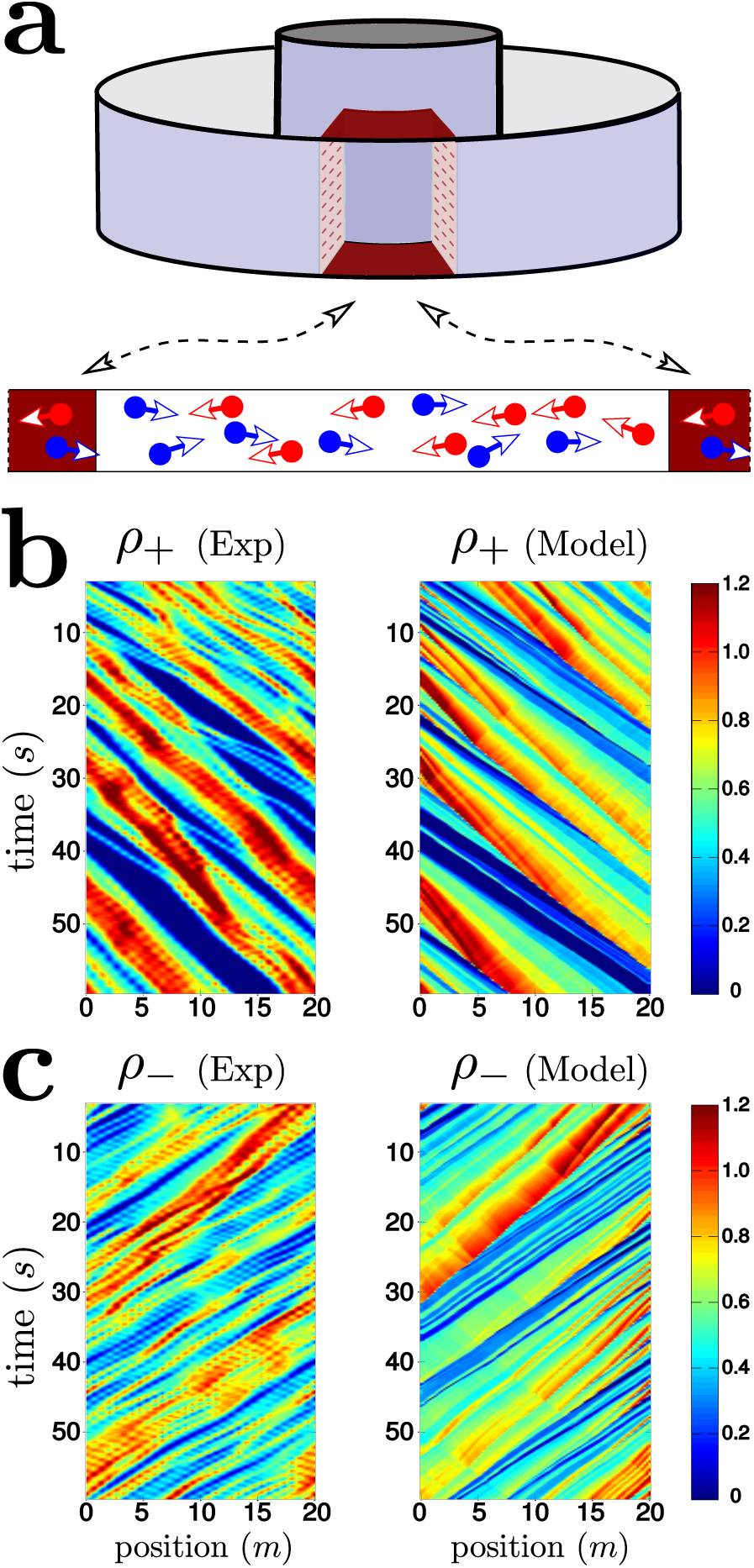
Model results and comparisons with experimental data. (**a**) Setting for the model: the density at the entry of the corridor (*x* = 0) is taken from the experiment and the model is used to predict the occupancy inside the corridor. (**b**-**c**) Clockwise and anti-clockwise (resp. **b** and **c**) pedestrian densities as functions of position (horizontal axis) and time (vertical axis running downwards), for one of the replications with 60 pedestrians corresponding to balanced fluxes (50% of pedestrians walking in each direction). Left picture of each panel: experiment; right picture: BM model, run with initial and boundary data and BFD estimated from the experimental data. The density is color-coded according to the lateral scale (in m^−2^).

**Figure 5:**
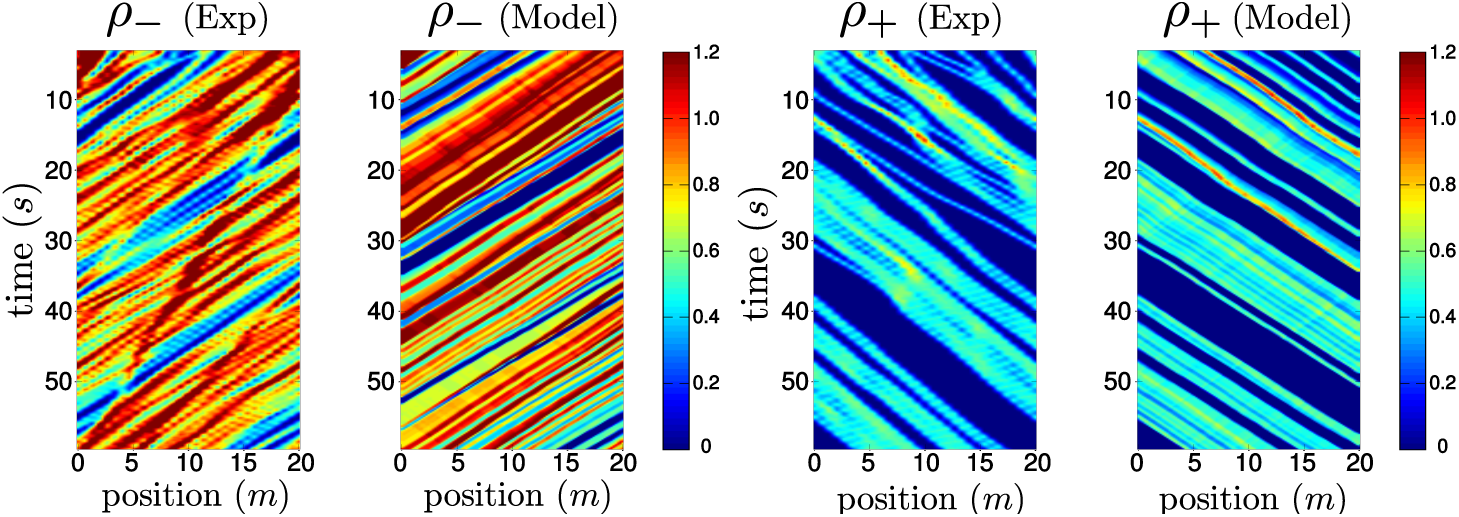
Model results and comparisons with experiment data with the 75%–25% flux balance: %75 are moving to the left (*ρ*_−_) and %25 are moving to the right (*ρ*_+_). See Fig. 4b,c for balanced flux.

### 3.3 Comparison of the model’s predictions with experimental data

To compare quantitatively the model with experimental data, we estimate the average and cluster velocities (resp. *u* and *λ*) defined in Eqs. 7 and 8. We detail in appendix B the numerical estimations of these curves. On Fig. 6, the difference *u – λ* between the average and cluster velocities is roughly linearly increasing. Using the expression of the flux (Eq. 1) together with the characteristics speeds (Eqs. 7 and 8), the theoretical prediction gives *u*_+_–*λ*_+_ ≈ 0.33 *ρ*_+_ ms^−1^ for small *ρ*_−_. According to this prediction, this offset should increase linearly with *ρ*_+_ and reach the value 0.4 ms^−1^ for *ρ*_+_ ≈ 1.2 m^−2^. This theoretical estimate provides the right order of magnitude of the experimentally observed value of *u*_+_–*λ*_+_.

**Figure 6:**
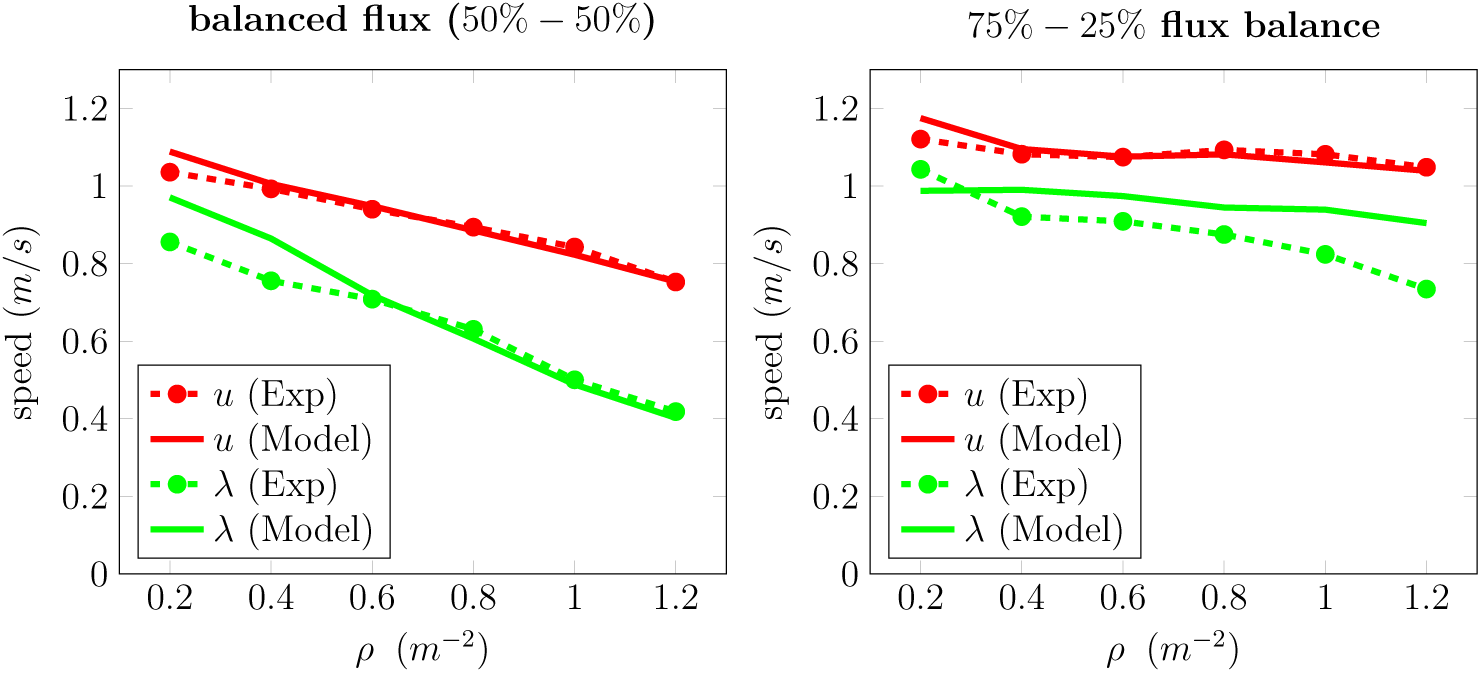
Pedestrian and cluster velocities. Comparison between the pedestrian velocity *u* (red curve) and cluster velocity *λ* (green curve) as functions of the local density of co-moving pedestrians *ρ* from the experiments (circles) and from the BM model (solid line) with balanced flux (50%–50%) on the left figure and with 75%–25% flux balance on the right figure. Since λ < *u*, information is propagating upstream as predicted by the BM model.

### 3.4 Segregation strategy efficiency

We now exploit the BM model to estimate the efficiency of a flow segregation strategy in which the two pedestrian types circulate preferably on one side of the corridor, with half the width of the corridor devoted to each type. In practice, this strategy can be achieved by suitable signaling. On the one hand segregation reduces the influence of counter-moving pedestrians, but on the other hand the available corridor width is smaller and the density of co-moving pedestrians increases. To measure the efficiency of the strategy, we test the model in the two situations. Let (*dN/dt*)_*S*_ and (*dN/dt*)_*NS*_ be the number of pedestrians per unit time that the corridor is able to accommodate in the segregated and non-segregated cases respectively, and *G* the gain in using the segregated strategy, i.e. the ratio of these two quantities minus one (see Methods). The gain as a function of the densities of the two pedestrian types is plotted in Fig. 7. It appears that the segregation strategy improves significantly the efficiency of the system when the densities of the two types are both large and of the same order. However, the strategy is inefficient when one type outnumbers the other one.In this case, it is more efficient to use the un-segregated strategy. This analysis demonstrates the benefits of a strategy optimization scheme in which the decision of implementing segregation or not could be taken in real-time. Indeed,segregation offers benefits only in some regions of the (*ρ*_+_*, ρ*_−_) plane. Since setting up a segregation strategy implies additional costs, the gain in traffic efficiency has to overtake the expense. The diagram presented in Fig. 7 helps finding which regimes of densities provide an overall benefit in separating the two inflows of pedestrians.

**Figure 7:**
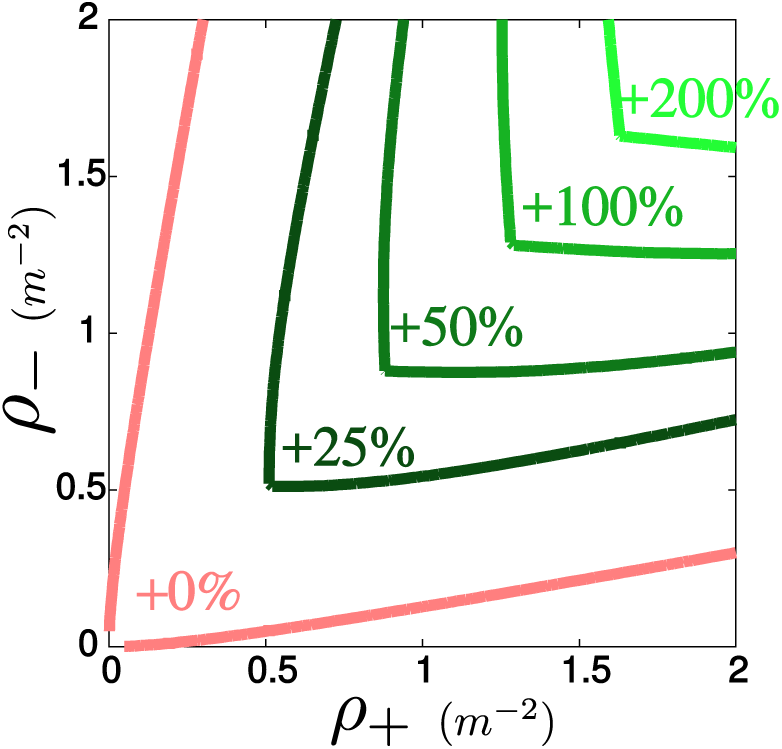
Efficiency segregation strategy. Estimation of the relative gain using the segregation strategy (in %) as a function of the densities *ρ*_+_*, ρ*_−_ (in m^−2^) of the two types of pedestrians (level curve representation). The strategy is efficient when both densities *ρ*_+_ and *ρ*_−_ are roughly balanced and large.

To estimate the efficiency of the segregation strategy, we compare the total flux of pedestrians when the two species of pedestrians *ρ*_+_ and *ρ*_−_ are mixed or segregated in acorridor. Without a segregation strategy, we suppose that the densities *ρ*_+_ and *ρ*_−_ areuniformly spread on the corridor. We assume here that the estimation of the flux *f* is independent as the corridor width. That will be more likely to be true for large corridor since the boundary effects would be less important. Thus, the number of pedestrians passing through a cross section per unit time is given by:

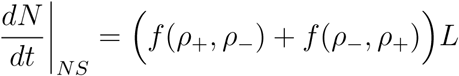

where *L* is the size of the corridor. The flux *f*_+_ is estimated from an interpolation of the three flux functions obtained in Table 1. More precisely, we use a quadratic regression to estimate each coefficient *a, b* and *c* at the ratio 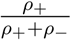 using the data of Tab. 1. Then,we use those coefficients to estimate *f*_+_(*ρ*_+_*, ρ*_−_) (see Eq. 9).

If the species *ρ*_+_ and *ρ*_−_ are segregated on each side of the corridor, the size of the corridor is divided by 2 and the density of co-moving pedestrian is multiplied by 2. Thus,we obtain:

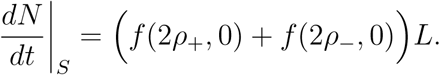

Then, the relative gain of the segregation strategy *G* is estimated through the following formula:

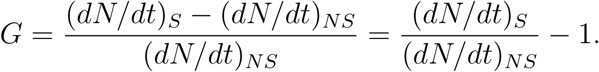

## 4 Discussion

In this work, we have set the objective to build simulation tools to assist the management of pedestrian traffic. Our attention was focused on the case of bidirectional flows because, on the one hand, it captures many common situations to be found in corridors or sidewalks, and on the other hand, it received relatively poor attention in the literature. One of our work hypothesis is that two main factors influence conditions in bidirectional traffic: density, and proportion of each directional flow.

In the first part of this work, we empirically analyze bidirectional traffic. Our main result is captured by Figure 3: we show an evidence of the influence of the density of each directional flow on the flux. To perform this empirical observation, we chose laboratory conditions. Choosing laboratory conditions allowed us controlling the goals and motivations of walkers, and limiting the effect of the other many factors which influence pedestrian behaviors, such as physiological (e.g., aging) or social ones (e.g., walking together in groups). We clearly isolate the role of the density factors we considered on the resulting traffic conditions. This however limited the amount of observation data due to the required effort to gather them (subjects recruitment, data reconstruction and processing efforts, etc.). Nevertheless, the presented method applies to any new data. As, for instance, the fundamental diagram may depend on the nature of the subjects that are using the walkway [35] (children, adults, aging people, etc.), it would be interesting to update the BFD from dataset with controlled population.

In the second part of this work, we introduce a first order model to estimate the bidirectional fundamental diagram. In spite of the relative simplicity of this model, we demonstrate the ability to adhere to observation data, as illustrated by our results reported in Table 1. We observe that the estimated average velocity *a* consistently takes similar values throughout the 3 sets of experiments. Friction with the co-moving pedestrians *b* is surprisingly larger in the experiments with a 50%–50% balance. It exceeds the friction with counter-moving pedestrians (i.e. *b > c*). One explanation of this result is the emergence of lanes. In the experiments with 50%–50% balance, lane formation is very strong whereas it is scarcely observed in the experiments with 75%–25% or 100%–0% balance. On the one hand, lane formation reduces the interactions with counter-moving pedestrians and thus decreases the friction coefficient *c*. But on the other hand, it enhances clustering with co-moving pedestrians and thus increases the friction coefficients *b*.

As a result, the friction coefficients *b* and *c* are associated with complex pattern formation in the longitudinal direction such as clustering as well as in the lateral direction such as lane formation. They provide a simple yet effective characterization of various flow regimes. The Bi-directional Macroscopic (BM) model critically depends on the three parameters *a, b* and *c*. By varying them, we can span various types of pedestrian behavior. As such, the BM model is capable of describing complex pedestrian behavior while having the efficiency of a one-dimensional model. Of course, one can make the flux expression *f* more complex to include all possible scenarios in a single expression. But this is at the cost of introducing new parameters which need to be estimated and this decreases the efficiency of the model in real-time crowd control situation. Therefore we prefer to use a *greedy modeling* approach with a simple (but not simplistic) model. The accuracy of the model can however be discussed with respect to the employed data-set. We have limited the role of secondary factors, we may expect to have a lower adherence using real, non-controlled, kinematics data (e.g., outdoor video tracking). This possible loss of accuracy need to be evaluated, nevertheless, we can expect that our results still hold in many cases, such as metro corridor traffic at rush, where most of the traffic is composed of pedestrian moving individually.

In the last part of this work, we have chosen to evaluate our model based on an analysis of cluster dynamics. The emergence of clusters as a consequence of density fluctuations is a classical example of spatiotemporal patterns in traffic flow phenomena [32, 33], also known as *Kinematic Waves* [34, 18]. Comparing the large-scale dynamical features of the clusters in the simulation and the data provides a practical way to assess the validity of the BM model. For a given type of pedestrians (clockwise or anti-clockwise), clusters are defined as regions of space where the density of this type of pedestrian exceeds a certain threshold value (see Methods). Fig. 6 gives the estimated average pedestrian velocity *u* (in red) and cluster velocity *λ* (in green) as functions of the density of pedestrians for the experiments (circles) and the simulations (solid lines) in the balanced flux case (50%–50%). We notice an excellent agreement between experiments and simulations despite some discrepancies at low density (i.e. *ρ*_+_ < .4*m*^−2^). Those discrepancies could be explained by the fluctuations in the experiments at low density which perturb the estimation of *u* and *λ*. Fig. 4 has already provided a qualitative comparison between traveling waves in the experiments and in the model, Fig. 6 provides a quantitative estimate of the resemblance. We notice that pedestrian velocity *u* is always faster than the cluster velocity *λ*. This indicates that the information is propagating upstream. Moreover, when the density increases, we observe a faster decay of the cluster velocity *λ* compared to the average velocity *u* (i.e. *u–λ* is increasing with *ρ*). This implies that perturbations propagate faster at large density as it is predicted by the BM model (see Methods).

## 5 Conclusion and Future Work

We have presented a study on bidirectional flows of pedestrian. We have first presented an experimental measurement of the bidirectional fundamental diagram, with changing conditions of flow proportions. We have then introduced a bidirectional traffic model, directly based on this diagram (a.k.a. “first order models “) for efficiency reasons. Finally, we demonstrate a relevant application of our model in the frame of automated traffic management systems, allowing users to estimate the benefit to flow segregation strategy to improve traffic.

The power of our approach is to capture complex phenomenon with a simple framework involving only three parameters. Moreover, one can easily extend the model to encompass more advanced features. As a first research direction, we would like to use real-time estimation of crowd parameters as it has been developed in several studies [36, 37, 38, 39, 40, 41, 42] to update the BFD diagram. The data assimilation strategy outlined above will be able to adapt the BM model quickly to changes in the nature and composition of the crowd.

Another possible extension would be to use models involving a differential relation between the flux and the local densities [43, 44] (a.k.a. “second order models”). Those models could generate metastable equilibria and phase transitions, which play an important role in traffic [45, 33]. Still, our first exploration of the data does not indicate that metastability plays an important role in pedestrian traffic.

To extend our work in two-dimensions (i.e. take into account the radial component), one has to consider the *direction* of the flow of pedestrians. Several macroscopic models have been proposed [46, 47, 48] with the introduction of a minimization principle to select the best route. Since most of bidirectional pedestrian flow have been studied at the microscopic level with Cellular Automata models [49, 50, 51, 52], it will be compelling to compare the two approaches.

Finally, it would be interesting to test the framework in a more complex environment such as a network of corridors. Several strategies could be tested to decongest traffic such as indicating which corridor to use depending on its occupancy. Hence, the efficiency of a network could be measured allowing to improve its design for safer evacuation.

## Appendix A. Analytic estimation of the cluster velocity λ

In a one-way traffic, there is only one type of density *ρ* and the flux is simply given by *f* (*ρ*). The characteristic velocities are then defined as (see Fig. 6b):

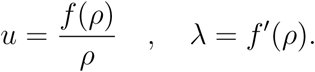

In a bi-directional traffic, there are two values for the cluster velocities, denoted by*λ*_*±*_(*ρ*_+_*,ρ*_−_). Indeed, the flux function *f* (*ρ*) must be replaced by the BFD *f* (*ρ*_+_*, ρ*_−_) and the quantity *f*′(*ρ*) must be replaced by the matrix

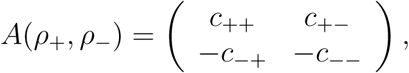

Where

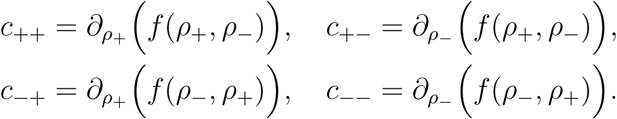

Cluster velocities *λ*_*±*_(*ρ*_+_*, ρ*_−_) are the eigenvalues of this matrix. They have been computed in [24] and are equal to

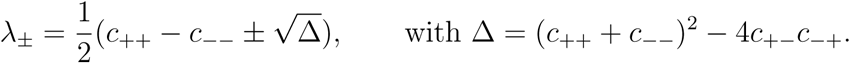

The eigenvalues *λ*_*±*_ are real (i.e. the BM model is hyperbolic [57]) if and only if Δ > 0. This is the case in the situation (met in our experiments) where

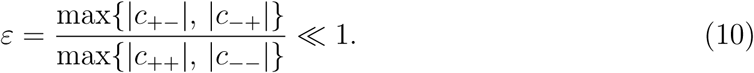

In general, it is not possible to associate one of these eigenvalues to either one or the other pedestrian species *ρ*_+_ and *ρ*_−_.For instance, if we denote *ρ*_1+_ and *ρ*_1−_ the perturbations of *ρ*_+_ and *ρ*_−_, there are not convected by *λ*_+_ and *λ*_−_. Instead, *λ*_+_ and *λ*_−_ are convecting the following linear combinations:

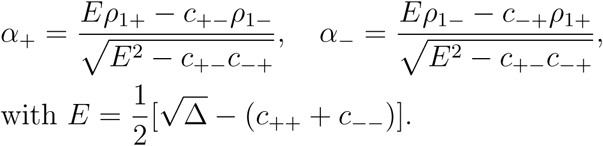

This means that the densities *ρ*_*±*_ are not purely convected. However, in the regime characterized by *ε* ≪ 1 (Eq. 10), we find that

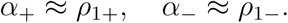

In this approximation, the densities *ρ*_*±*_ are purely convected with velocities *λ*_*±*_, which means that we can associate *λ*_+_ and *λ*_−_ to the velocities of clusters of the clockwise and anti-clockwise moving pedestrians respectively. In this approximation, to the first order in *ε*, we find:

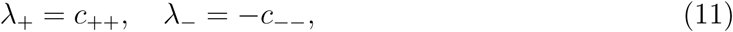

which gives a situation similar to car traffic.

## Appendix B. Identification and quantification of clusters

Cluster analysis provides a lens to analyze emergence of macroscopic structures in pedestrians dynamics. From a macroscopic viewpoint, a cluster at a given time *t* is a connected set of points *x* where the density *ρ*(*x, t*) is larger than a given threshold *h* (see Fig. 8a). Obviously, there are different clusters for the clockwise (*ρ* = *ρ*_+_) and anticlockwise (*ρ* = *ρ*_−_) pedestrians. As clusters move in time, they are bounded by level curves *X*(*t*) of *ρ* defined by:

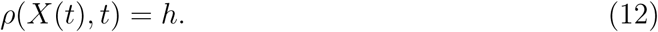

**Figure 8:**
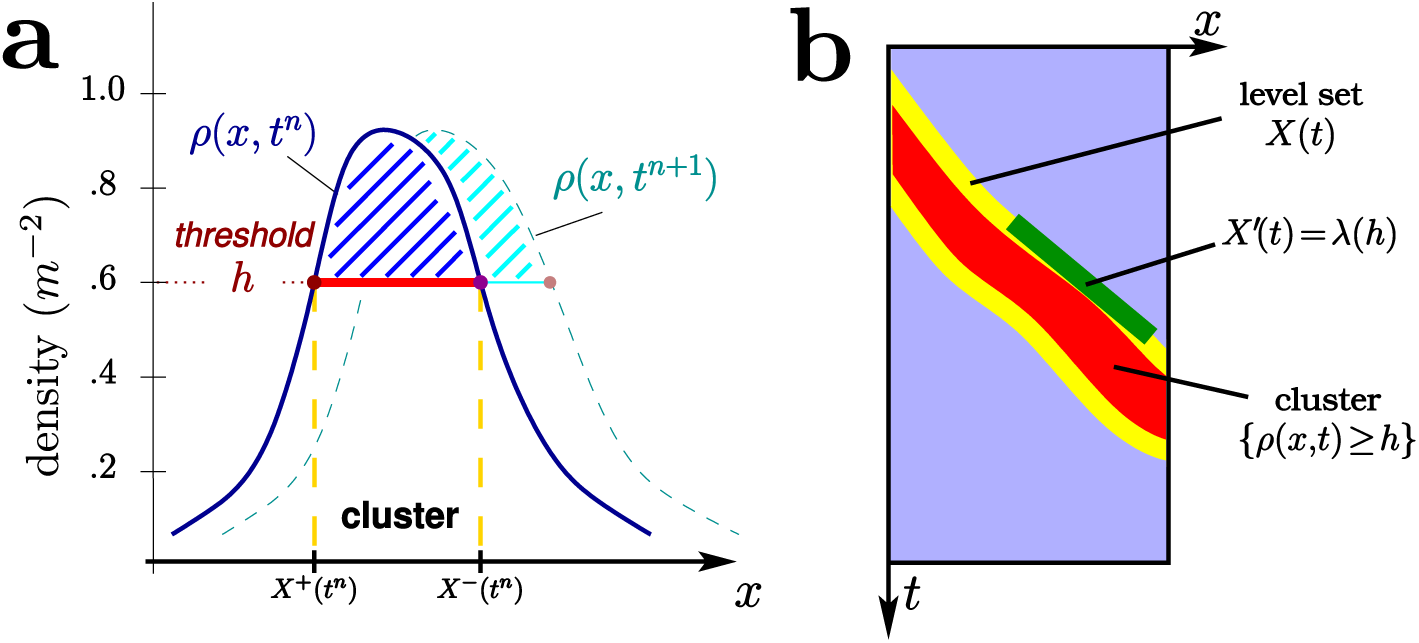
Cluster and cluster velocity. (**a**) Illustration of a cluster defined from a density distribution *ρ*(*x, t*^*n*^) for a given threshold *h*. The edges of the cluster (i.e. *X*^+^ and *X*^−^) described the level curves of *ρ*(*x, t*). (**b**) Graphical representation of the clustervelocity using the level curves of *ρ*.The picture depicts the edges (in yellow) of the cluster(in red). The cluster edge velocity i.e. the slope of the cluster edge in the position-time plane is given by *f*′(*h*) and is illustrated by the green segment.

To determine a cluster, we first need to construct the cluster boundaries, i.e. the curves *X*(*t*) defined by (12), from the pointwise estimation of *ρ*_*±*_.

To construct such curves *X*(*t*), we consider *ρ*^*n*^(*x*) the continuous piecewise linearn function of *x* which interpolates the estimated density 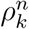 at a given time *t*_*n*_ (here, we indifferently use the notation *ρ* for *ρ*_+_ and *ρ*_−_; the ± indices will be used below to designate the start and end points of the clusters). For a given threshold *h*, we determine two sets of points 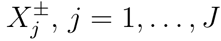 such that 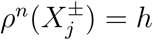 and 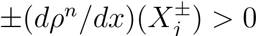. The number of points in each set is the same, *J*, due to the periodicity of *ρ*^*n*^.It is finite unless *ρ*^*n*^(*x*) has constant value *h* on a whole interval, a nongeneric situation that we discard.

The *j*-th cluster is defined as the interval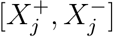, because, due to the conditions on the derivatives, we have *ρ*^*n*^(*x*) ≥ *h* for all *x* belonging to this interval (see Fig. 8). We now consider two consecutive time intervals *t*^*n*^ and *t*^*n*^+^1^. We connect 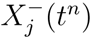 to 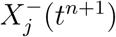 and 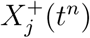 to 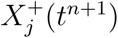, if the two consecutive points are closer than a threshold value set to *λ*_0_Δ*t*, with *λ*_0_ = 3.3 ms^−1^. The temporal sequences of connected points 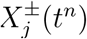 define, after time-interpolation between the discrete time values, some level curves 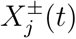. When the matching procedure defined above finds no solution, the corresponding level curve is ended.

This procedure can be repeated for different levels *h*, and yields different families of level curves *X*(*t*) (see Fig. 9a). When the time interval of existence of a level curve (or lifetime) is too short, that level curve is considered as irrelevant and is removed. Minimal existence time of 1, 2 and 3 seconds have been retained (see Fig. 9b for the example of a cutoff at 3 seconds). In order to remove further oscillations of the level curves (which are associated to the spatial discretization of Δ*x* = 0.6 m), we filter them by convolution in time with a truncated Gaussian function with a support larger than Δ*x*, adapting the normalization of the Gaussian to accommodate for the beginning and the end of the level curve (see Fig. 9c).

**Figure 9:**
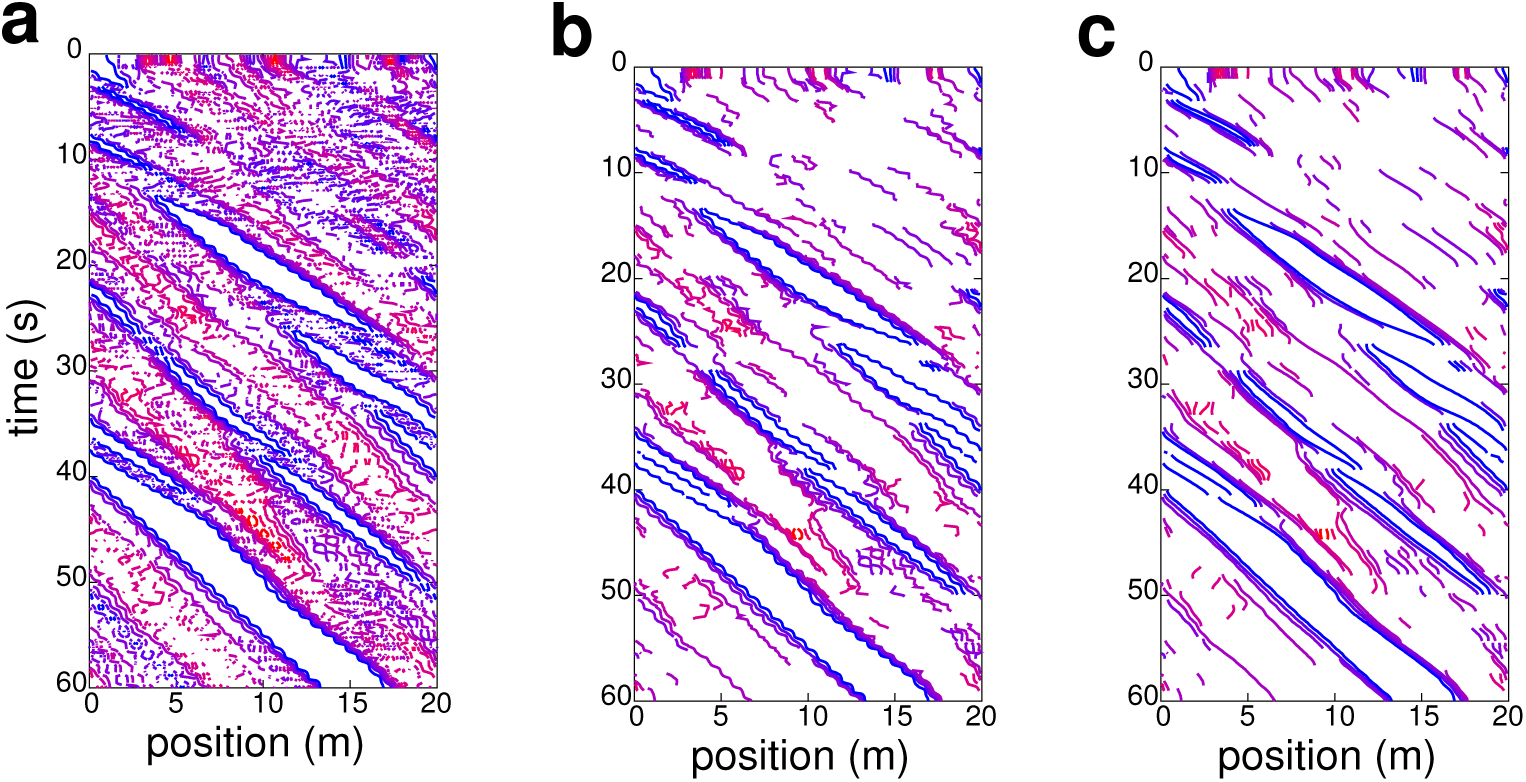
Level curves of the density. Level curves *X*(*t*) for various levels *h* for the replication displayed in Fig. 4 and clockwise pedestrian density *ρ*_+_. Horizontal axis is space and vertical axis is time, running downwards. The color code corresponds to the level height *h*, from blue (lower levels) to red (higher levels). (**a**): with no cutoff and no filtering. (**b**): after cutoff (only level curves with life-time greater than 3 seconds are kept) but before filtering. (**c**): after cutoff and filtering.

From the level curves *X*(*t*), we can then deduce the cluster velocity *λ*, that is the velocity at which the level curves are transported (see Fig. 8b). One should rather speak of ‚cluster edge velocity’ since clusters have two boundaries, each of them having its own speed, but as they are of the same order, we will use the term ‘cluster velocity‚ for short. To estimate the cluster velocity *λ* on each level curve of density level *h*, we compute its average slope. The slopes of the different connected components of a given level curve are computed independently and later on averaged. A weighted average with weights equal to the lifetime of the corresponding connected component is applied. This procedure leads to an estimate of *λ* as a function of the specific density level *h*. Thus, if we restore the indication *ρ* for the density level *h*, we obtain *λ* as a function of *ρ*.

## Acknowledgment

This work has been supported by the French ‚Agence Nationale pour la Recherche (ANR)‚ in the frame of the contracts ‚Pedigree‚ (ANR-08-SYSC-015-01) and ‚CBDif-Fr‚ (ANR-08-BLAN-0333-01). PD acknowledges support by the Engineering and Physical Sciences Research Council (EPSRC) under grant no. EP/M006883/1, by the Royal Society and the Wolfson Foundation through a Royal Society Wolfson Research Merit Award no. WM130048 and by the National Science Foundation (NSF) under grant no. RNMS11-07444 (KI-Net). PD is on leave from CNRS, Institut de Mathematiques de Toulouse, France SM was supported in part by NSF grants #1107444 (KI-Net) and #1515592.

## Data accessibility

Data supporting this work are available on https://figshare.com/articles/README/4056495 and https://figshare.com/articles/Experimentaldata/4056483

## References

[1] Martine G, Marshall A. 2007 State of world population 2007: unleashing the potential of urban growth. UNFPA.

[2] Rietveld P. 2000 Non-motorised modes in transport systems: a multimodal chain perspective for The Netherlands. Transportation Research Part D: Transport and Environment 5(1), 31–36.

[3] Papageorgiou M., Diakaki C., Dinopoulou V., Kotsialos A. and Wang Y. 2003 Review of road traffic control strategies. Proceedings of the IEEE 91(12), 2043–2067

[4] Still, G.K. 2000 Crowd Dynamics. PhD Thesis, University of Warwick

[5] Helbing, D. et al. 2005 Self-organized pedestrian crowd dynamics: Experiments, simulations, and design solutions Transport Sci 39, 1–24.

[6] Shiwakoti N, Sarvi M. 2013 Enhancing the panic escape of crowd through architectural design Transportation Research Part C: Emerging Technologies 37, 260–267.

[7] Benfold B., Reid I. 2011 Stable multi-target tracking in real-time surveillance video Computer Vision and Pattern Recognition (CVPR), 2011 IEEE Conference on 3457– 3464.

[8] Kratz, L. and Nishino, K. 2012 Tracking Pedestrians using Local Spatio-Temporal Motion Patterns in Extremely Crowded Scenes IEEE Trans. on Pattern Analysis and Machine Intelligence 34 987–1002.

[9] Plaue M, Chen M, Bärwolff G, Schwandt H. 2011 Trajectory extraction and density analysis of intersecting pedestrian flows from video recordings. Photogrammetric Image Analysis, Springer, 285–296.

[10] Ali, S. and Shah, M. 2008 Floor Fields for Tracking in High Density Crowd Scenes Computer Vision – ECCV 2008: 10th European Conference on Computer Vision 1–14.

[11] Zhang J, Klingsch W., Schadschneider A, Seyfried A. 2012 Ordering in bidirectional pedestrian flows and its influence on the fundamental diagram. J. Stat. Mech. Theory Exp. 02, P02002.

[12] Saberi M, Aghabayk K, Sobhani A. 2016 Spatial fluctuations of pedestrian velocities in bidirectional streams: Exploring the effects of self-organization. Physica A: Statistical Mechanics and its Applications. 434, 120–128.

[13] Helbing, D. and Molnar, P. 1995 Social Force Model for Pedestrian Dynamics. Physical Review E. 51 4282–4286.

[14] Reynolds, C. W 1987 Flocks, herds and schools: A distributed behavioral model ACM SIGGRAPH Computer Graphics 21 25–34.

[15] Ondrej, J. et al. 2010 A synthetic-vision based steering approach for crowd simulation ACM Transactions on Graphics 29, 1–9.

[16] Moussaid, M. et al. 2011 How simple rules determine pedestrian behavior and crowd disasters Proc Natl Acad Sci. 108, 6884–6888.

[17] Lemercier S, et al. 2011 Reconstructing motion capture data for human crowd study. Motion in Games, 365–376.

[18] Lemercier S et al. 2012 Realistic following behaviors for crowd simulation. Comput Graph Forum. 31, 489–498.

[19] Rosini, M. 2013 Macroscopic Models for Vehicular Flows and Crowd Dynamics: Theory and Applications Springer.

[20] Lighthill M J, Whitham G B. 1955 On kinematic waves. II. A theory of traffic flow on long crowded roads. P Roy Soc Lond A Mat. 229, 317–345.

[21] AlGadhi S A, Mahmassani H. S., Herman R. 2002 A speed-concentration relation for bi-directional crowd movements with strong interaction. Pedestrian and evacuation dynamics. 3–20.

[22] Kretz T et al. 2006 Experimental study of pedestrian counterflow in a corridor. J Stat Mech-Theory E. 2006, P10001.

[23] Lam W H et al. 2003 A generalised function for modeling bi-directional flow effects on indoor walkways in Hong Kong. Transport Res A-Pol. 37, 789–810.

[24] Appert-Rolland C, Degond P, Motsch S. 2011. Two-way multi-lane traffic model for pedestrians in corridors. Netw Heterog Media. 6, 351–381

[25] Tory E, et al. 2011 An adaptive finite-volume method for a model of two-phase pedestrian flow. Netw Heterog Media. 6, 401–423

[26] Bick J H, Newell G F. 1960 A continuum model for two-directional traffic flow. Q. Appl. Math.

[27] Goatin P, Mimault M. 2014 A mixed system modeling two-directional pedestrian flows. Math Biosci Eng. 12, 375–392

[28] Helbing D et al. 2015 Saving human lives: what complexity science and information systems can contribute. J Stat Phys. 158, 735–781

[29] Helbing D, Mukerji P. 2012 Crowd disasters as systemic failures: analysis of the Love Parade disaster. EPJ Data Science. 1, 1–40.

[30] Moussaid M, et al. 2012 Traffic instabilities in self-organized pedestrian crowds. Plos Comput Biol. 8, e1002442.

[31] Kirchner A, Nishinari K, Schadschneider A. 2003 Friction effects and clogging in a cellular automaton model for pedestrian dynamics. Phys Rev E. 67, 056122.

[32] Helbing D. 2001. Traffic and related self-driven many-particle systems. Rev Mod Phys.73, 1067.

[33] Kerner B S. 2004 The physics of traffic: empirical freeway pattern features, engineering applications, and theory. Springer Verlag.

[34] Lighthill M J, Whitham G B. 1955 On kinematic waves. I. Flood movement in long rivers. Proc R Soc Lon Ser-A. 229, 281–316.

[35] Henderson L F. 1971 The statistics of crowd fluids. Nature. 229, 381–383.

[36] Marana A N et al. 2005 Real-time crowd density estimation using images. Lect Notes Comput Sc. 355–362.

[37] Zhou B, Zhang F, Peng L. 2012 Higher-order SVD analysis for crowd density estimation. Comput Vis Image Und. 116, 1014–1021.

[38] Yaseen S et al. 2013 Real-time crowd density mapping using a novel sensory fusion model of infrared and visual systems. Safety Sci. 57, 313–325.

[39] Johansson A et al. 2008 From crowd dynamics to crowd safety: a video-based analysis. Adv Complex Syst. 11, 497–527.

[40] Helbing D, Johansson A, Al-Abideen H Z. 2007 Dynamics of crowd disasters: An empirical study. Phys Rev E. 75, 046109.

[41] Wirz M et al. 2013 Probing crowd density through smartphones in city-scale mass gatherings. EPJ Data Science. 2, 1–24.

[42] Feliciani C, Nishinari K. 2016 Empirical analysis of the lane formation process in bidirectional pedestrian flow. Physical Review E. 94(3), 032304.

[43] Aw A, Rascle M. 2000 Resurrection of “second order” models of traffic flow. SIAM J Appl Math. 60, 916–938.

[44] Payne H J. 1979 FREFLO: A macroscopic simulation model of freeway traffic. Transp Res Record.

[45] Klar A et al. 2004 Multivalued Fundamental Diagrams and Stop and Go Waves for Continuum Traffic Flow Equations. SIAM J Appl Math. 64, 468–483.

[46] Hughes R L. 2002 A continuum theory for the flow of pedestrians. Transport Res B-Meth. 36, 507–535.

[47] Hoogendoorn S P, Bovy P H. 2004 Pedestrian route-choice and activity scheduling theory and models. Transport Res B-Meth. 38, 169–190.

[48] Bellomo N, Piccoli B, Tosin A. 2012 Modeling crowd dynamics from a complex system viewpoint. Math Mod Meth Appl S. 22, 1230004.

[49] Blue V J, Adler J L. 1999 Cellular automata microsimulation of bidirectional pedestrian flows. Trans Res B. 1678, 135–141.

[50] Jian L, Lizhong Y, Daoliang Z. 2005 Simulation of bi-direction pedestrian movement in corridor. Physica A. 354, 619–628.

[51] Weng W G et al. 2007 A behavior-based model for pedestrian counter flow. Physica A. 375, 668–678.

[52] Flötteröd G., Lämmel, G. 2015 Bidirectional pedestrian fundamental diagram. Transportation Research Part B: Methodological. 71, 194–212.

[53] Daamen W, Hoogendoorn S P. 2003 Controlled experiments to derive walking behaviour. Eur J of Trans Infrastruct Res. 3, 39–59.

[54] Birdsall C K, Langdon A B. 2014 Plasma physics via computer simulation. CRC Press.

[55] Fehske H, Schneider R, Weisse A. 2007 Computational Many-Particle Physics. Springer.

[56] LeVeque R J. 2002 Finite volume methods for hyperbolic problems. Cambridge University Press.

[57] LeVeque R J. 1992 Numerical Methods for Conservation Laws. Birkhäuser.

[58] Kurganov A, Tadmor E. 2000 New high-resolution central schemes for nonlinear conservation laws and convection-diffusion equations. J Comput Phys. 160, 241–282.

